# Rationalize the Functional Roles of Protein-Protein Interactions in Targeted Protein Degradation by Kinetic Monte-Carlo Simulations

**DOI:** 10.1101/2024.09.26.615190

**Authors:** Zhaoqian Su, Shanye Yin, Yinghao Wu

## Abstract

Targeted protein degradation is a promising therapeutic strategy to tackle disease-causing proteins that lack binding pockets for traditional small-molecule inhibitors. Its first step is to trigger the proximity between a ubiquitin ligase complex and a target protein through a heterobifunctional molecule, such as proteolysis targeting chimeras (PROTACs), leading to the formation of a ternary complex. The properties of protein-protein interactions play an important regulatory role during this process, which can be reflected by binding cooperativity. Unfortunately, although computer-aided drug design has become a cornerstone of modern drug development, the endeavor to model targeted protein degradation is still in its infancy. The development of computational tools to understand the impacts of protein-protein interactions on targeted protein degradation, therefore, is highly demanded. To reach this goal, we constructed a non-redundant structural benchmark of the most updated ternary complexes and applied a kinetic Monte-Carlo method to simulate the association between ligases and PROTAC-targeted proteins in the benchmark. Our results show that proteins in most complexes with positive cooperativity tend to associate into native-like configurations more often. In contrast, proteins very likely failed to associate into native-like configurations in complexes with negative cooperativity. Moreover, we compared the protein-protein association through different interfaces generated from molecular docking. The native-like binding interface shows a higher association probability than all the other alternative interfaces only in the complex with positive cooperativity. These observations support the idea that the formation of ternary complexes is closely regulated by the binary interactions between proteins. Finally, we applied our method to cyclin-dependent kinases 4 and 6 (CDK4/6). We found that their interactions with the ligase are not as similar as their structures. Altogether, our study paves the way for understanding the role of protein-protein interactions in PROTACE-induced ternary complex formation. It can potentially help in searching for degraders that selectively target specific proteins.

## Introduction

After decades of progress in drug discovery, there are still up to 90% of human proteins considered to be ‘undruggable’ [1]. This conclusion is based on the traditional theory of inhibitor design used to block the active sites on the surfaces of proteins [2]. Recent efforts have been made by different strategies to target these “undruggable” proteins [3], including covalent regulation [4], allosteric regulation [5], and antibody-drug conjugates [6]. Among these strategies, targeted protein degradation (TPD) has drawn tremendous attention due to its higher potency and lower risk of off-target effects compared to traditional small-molecule inhibitors [7]. Proteolysis targeting chimeras (PROTACs), as one of the most well-known examples of TPD, are heterobifunctional molecules [8]. They contain two small-molecule moieties with a linker to induce proximity between a ubiquitin ligase complex and a target protein, leading to its ubiquitination and degradation [9]. As a result, proteins that lack an active site can be targeted. Given this potential for expanding the druggable space, a large variety of PROTACs are currently in clinical trials [10, 11].

The connection between a target protein and the E3 ligase by PROTACs results in the formation of a ternary complex [12]. Several features are used to characterize the productivity of PROTAC-induced ternary complexes [13]. One important feature is cooperativity (α) [14]. It is defined as dividing a PROTAC’s binary dissociation constant by its ternary dissociation constant [15], implying that the protein-small molecule interactions can be influenced by the protein-protein interactions (PPIs) in the ternary complex. A positive cooperativity (α>1) indicates that the association between the target protein and the ligase facilitates ternary complex formation due to the energetically favored interactions obtained at the interface between two proteins. On the other hand, a positive cooperativity (α<1) suggests that the PPI is unfavorable to the ternary complex formation for reasons such as steric clashes. It has been found that positive cooperativities can ensure selectivity and elicit potent degradation, highlighting the important role of PPIs in improving the degradation efficiency of PROTACs.

The ternary complex formation can be measured by fluorescence polarization, isothermal titration calorimetry (ITC) or time-resolved fluorescence energy transfer (TR-FRET) [16]. Because these *in vitro* approaches do not adequately capture the complexity of the cellular environment, additional methods such as NanoBRET and NanoBiT are used to detect the formation of ternary complexes in intact cells [17]. However, they are not sensitive to the weak protein-protein interactions in the ternary complex. In addition to the experimental technologies, the computational model has become a cornerstone of modern drug development [18–20]. Various docking-based pipelines, such as PRosettaC [21], PatchDock [22], and FRDOCK [23], have recently been established specifically for TPD. Along with the accumulation of degradation data in the literature, online databases, such as PROTAC-DB [24], have been constructed, which enable the implementation of machine learning strategies to assess the properties of PROTACs, including degradability and permeability [25]. However, compared to the traditional inhibitor design, the power of these computational approaches is limited by the small number of ternary complexes that are currently available in the Protein Data Bank (PDB). Moreover, most of these approaches focus on modeling the final structures of ternary complexes. Very few of them can be used to study the process of PROTAC-induced ternary complex formation. As a result, they are not able to estimate the impact of cooperativity and understand the function of PPIs in targeting PROTACs.

We have previously developed a kinetic Monte-Carlo (KMC) algorithm to simulate the association between two proteins guided by a physics-based scoring function [26]. A strong correlation was obtained between our calculations and the experimentally measured association rates. Here, we employed the method to the complexes formed by the E3 ligase and PROTAC-targeted proteins. We collected the structures for a non-redundant set of the most updated ternary complexes from the PDB. By testing our method on the dataset, we found that proteins in most complexes with positive cooperativity tend to associate into native-like configurations more often. In contrast, proteins very likely failed to associate into native-like configurations in complexes with a negative cooperativity. Moreover, for the complex with high cooperativity, the probability of protein-protein association through the native-like interface is much higher than those through the decoy interfaces generated from protein-protein docking methods. These results suggest that the association probability derived from our simulations could be a strong indicator for the cooperativity of ternary complex formation. Finally, we applied our KMC simulation to a case study of TPD for cyclin-dependent kinases 4 and 6 (CDK4/6) [27]. Although these two proteins share high structural similarity, we found that their associations with the ligase are quite different. This observation could explain why the same PROTAC can cause selective degradation between CDK4 and CDK6. In summary, our method provided insights to the molecular mechanism of PROTACE-induced ternary complex formation. It will be useful to search for the optimal protein-protein binding interfaces in order to design PROTACs with higher potency and specificity.

## Results

### Proteins are easier to associate together in the ternary complexes formed with positive cooperativity

Protein structures are essential to carrying out their functions. The formation of a ternary complex is the crucial step of targeted degradation. The crystallographic structures of ternary complexes for various systems have recently been deposited in the PDB. We constructed a dataset of PROTAC-based ternary complexes. The dataset contains 9 complexes formed by different target proteins and ligases. The atomic structures of these complexes have all been experimentally determined. The detailed process of collecting these complexes is described in the **Methods**. More information about the dataset can be found in **Table 1**. **Figure 1** plots the structures of all 9 ternary complexes in our dataset. The target proteins in these complexes are shown in red and the ligases are shown in green. The PROTACs between the ligases and target proteins are highlighted by the Van Der Waals representation. The corresponding PDB id is indicated below each complex. The experimentally measured values of cooperativity have also been obtained from the literature for all complexes.

**Figure 1:**
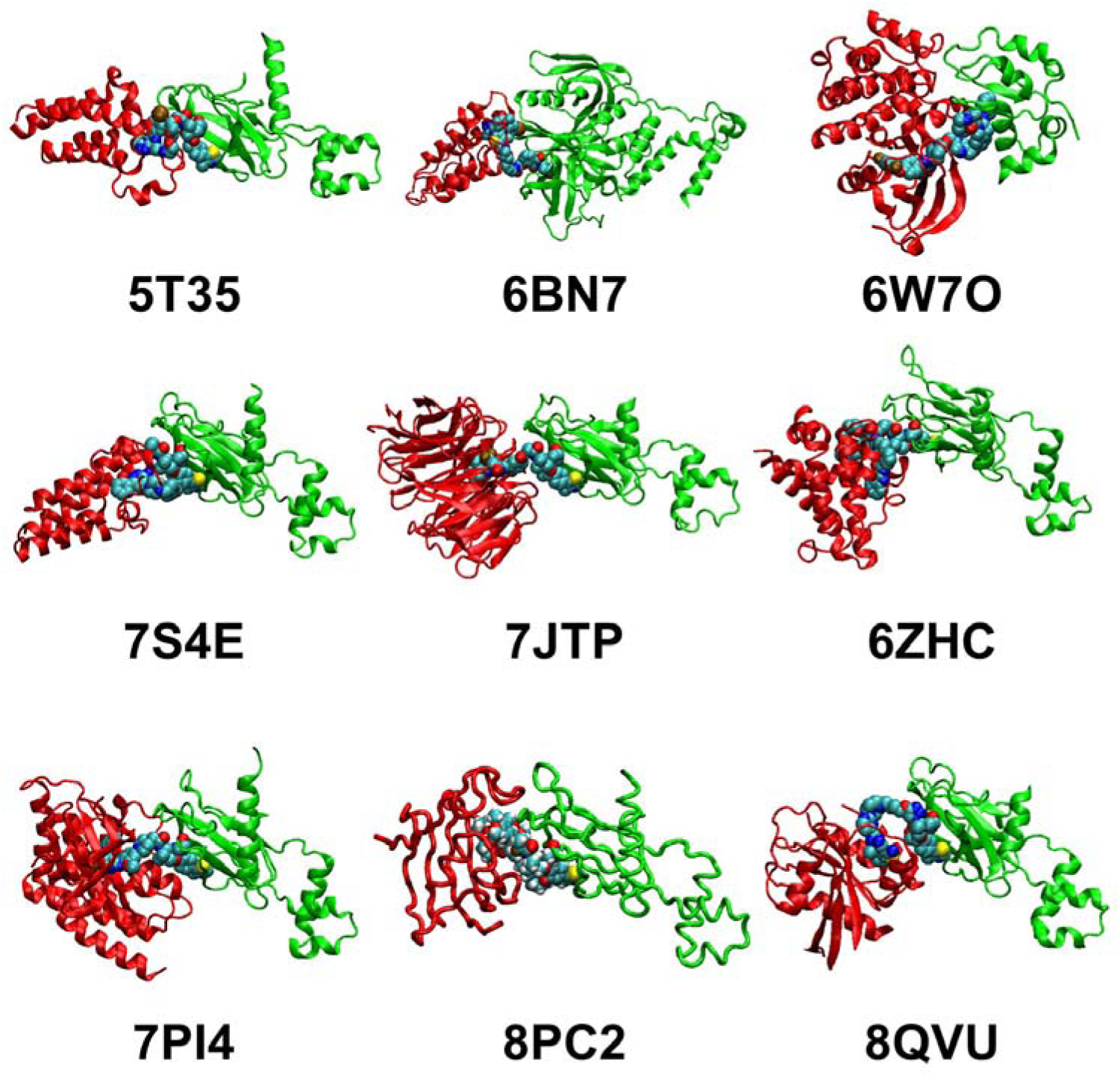
We constructed a benchmark dataset consisting of 9 PROTAC-based ternary complexes. This non-redundant dataset contains interactions between different types of target proteins and ligases. The structures of these complexes were downloaded from the PDB, and are shown in this figure. The target proteins in these complexes are plotted in red with cartoon representation. The ligases in these complexes are shown in green with cartoon representation. The PROTACs between the ligases and target proteins are highlighted by the Van Der Waals representation. The corresponding PDB id of each complex is indicated below its corresponding structure.

**Table 1.**

| PDB ID | Target Protein | Ligase | Association Probability | Cooperativity | References |
| --- | --- | --- | --- | --- | --- |
| 5T35 | BRD4<sup>BD2</sup> | VHL | 0.338 | + | [35] |
| 6BN7 | BRD4<sup>BD1</sup> | CRBN | 0.042 | - | [13] |
| 6W7O | BTK | cIAP | 0.051 | - | [36] |
| 7S4E | SMARCA2 | VHL | 0.104 | + | [37] |
| 7JTP | WDR5 | VHL | 0.173 | + | [38] |
| 6ZHC | BCL2L | VHL | 0.037 | - | [39] |
| 7PI4 | FAK | VHL | 0.0 | + | [40] |
| 8PC2 | FKBP5 | VHL | 0.0 | - | [41] |
| 8QVU | KRAS | VHL | 0.162 | + | [42] |

For all nine ternary complexes, we applied our KMC to simulate the process of protein-protein association. We downloaded the atomic structures of all complexes from the PDB. Because this study focuses on the function of interactions between proteins in ternary complexes, the PROTACs were not included in our simulations. The detailed algorithm of the KMC simulation is described in the **Methods**. For each complex, the KMC generated 10^3^ simulation trajectories from different initial configurations. Among these trajectories, some can successfully form native-like conformations of a protein-protein complex at the end of simulations, while others failed to associate together. Using the complex formed by the target protein BRD4^BD2^ and the ligase VHL (PDB id 5T35) as an example, the KMC results of forming a native-like interaction and not forming an interaction are plotted in **Figures 2a** and **Figure 2b**, respectively. Moreover, for a specific complex, we counted the number of trajectories in which native-like complexes were observed, and further calculated the association probability of the complex by dividing the number over all the 10^3^ trajectories. As a result, the calculated association probabilities for all complexes in the dataset are listed in **Table 1**.

**Figure 2:**
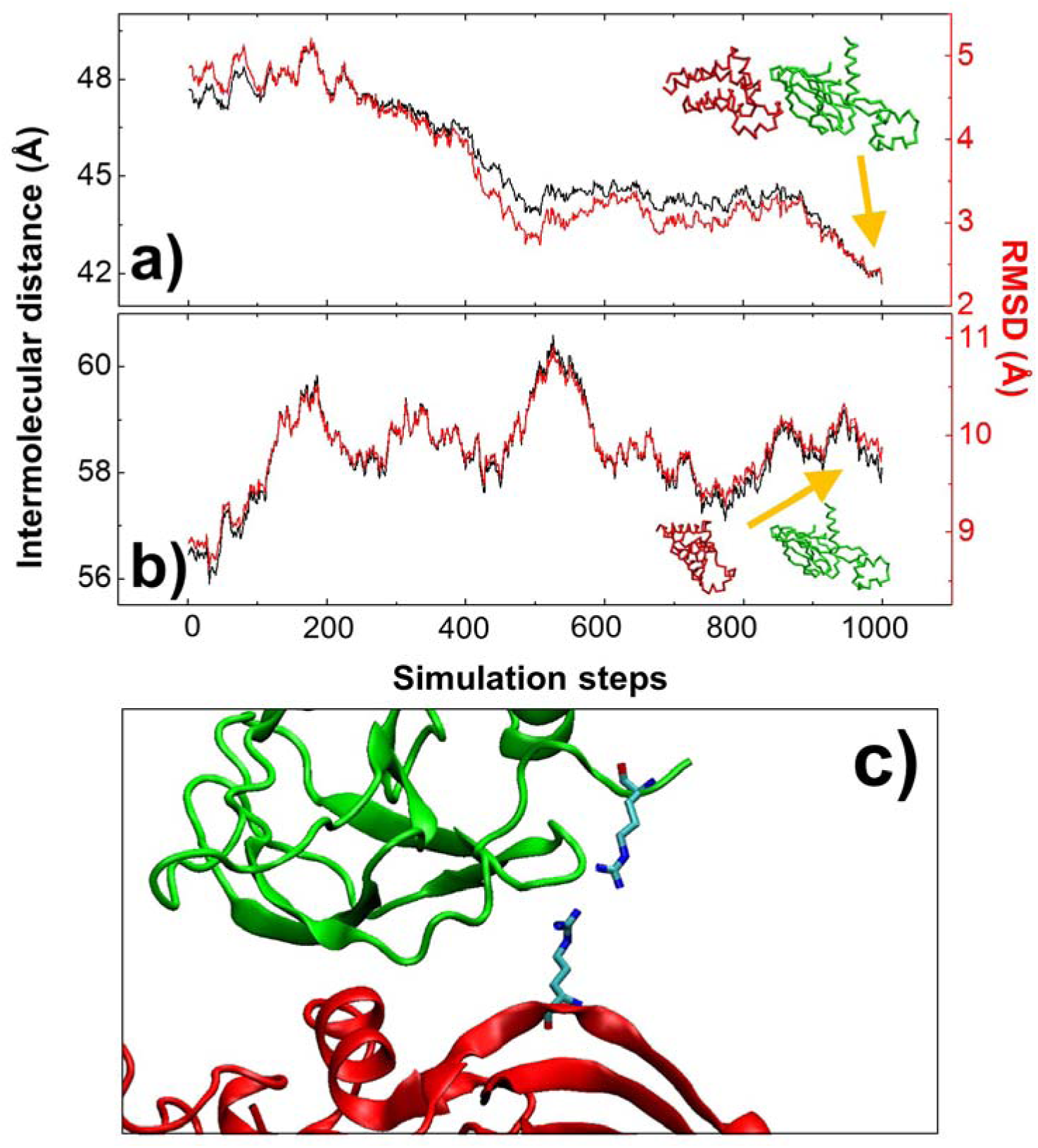
We carried out 10^3^ KMC simulation trajectories to each complex in the benchmark dataset. Two representative trajectories are plotted to illustrate how the distance between the centers of mass for the two monomers (black curves) and the root-mean-squared-distance (RMSD) from the native complex (red curves) changed with the simulation time. In **(a)**, both intermolecular distance and RMSD decrease with the simulation time, indicating two proteins successfully associated into a native-like complex, as shown by the inserted panel in the figure. In **(b)**, on the other hand, both intermolecular distance and RMSD increase throughout the end of the simulation, indicating two proteins failed to associate, as shown by the inserted panel in the figure. As the only exception of our KMC simulations, no native-like conformation was found for a ternary complex (PDB id 7PI4) with measured positive binding cooperativity. In a close-up view at the interface of this complex **(c)**, we found an Arginine from the target protein (red) forming a contact with another Argine from the ligase (green). This repulsive interaction led to the result that two proteins could not effectively associate together in our KMC simulations.

Our simulation results indicate a strong correlation between the calculated association probability and cooperativity. In the dataset, five out of nine complexes show positive cooperativity, while the remaining four complexes show negative cooperativity. For the five complexes with positive cooperativity, our calculated association probabilities are all higher than 0.1 except 7PI4 (**Table 1**). On the other hand, the calculated association probabilities for the four complexes with negative cooperativity are all lower than 0.1. We applied a student’s t-test to assess the statistical significance of our observed difference in association probability between the ternary complexes with positive cooperativity and the ternary complexes with negative cooperativity. The average association probability of complex with positive cooperativity (except 7PI4) is 0.19 and the standard deviation of the distribution is 0.11, while the average association probability of complex with negative cooperativity is 0.03 and the standard deviation is 0.02. The null hypothesis is that there is no statistical difference between two groups. The hypothesis was tested at a 95% confidence interval, and the calculated t-score equals 3.14 and the corresponding P-value is 0.02. The small P-value for the t-test suggests that the null hypothesis can be rejected and the alternative hypothesis, i.e., the association probability of complexes with positive cooperativity are significantly higher than the association probability of complexes with negative cooperativity, should be accepted.

The only exception of our KMC simulation is the complex formed between target protein FAK and ligase VHL (PDB id 7PI4). The experimental evidence showed that the ternary complex was formed by a positive cooperativity. However, no native-like conformation was found among the 10^3^ trajectories generated by KMC simulations. As a result, the calculated association probability of 7PI4 is 0. In order to understand why FAK and VHL could not form interactions in our simulation, we plotted a close-up view at the interface between the two proteins (**Figure 2c**). Interestingly, the figure shows that an Arginine from FAK (red) forms a contact with another Argine from VHL (green). The interaction between the two positively charged sidechains is repulsive and thus energetically unfavored. We speculate that the energetically unfavored interface between FAK and VHL prevents them from forming a complex in our simulations. It is likely that an alternative energetically more favored binding interface exists in the presence of the PROTAC molecule, leading to positive cooperativity as experimentally measured.

Taken together, we tested our computational simulations on a comprehensive dataset consisting of various target proteins and ligases. We found that proteins in the ternary complexes with positive cooperativity are more likely to interact with each other. In contrast, proteins in the ternary complexes with negative cooperativity are less likely to interact with each other. This result suggests that the formation of ternary complexes is closely regulated by the binary interactions between proteins.

### The protein-protein binding interfaces in the ternary complex with high cooperativity are preferred to other potential interfaces

The structures of ternary complexes in the PDB are determined with the presence of PROTACs. Previous studies showed that proteins can interact with each other through multiple conformations during the formation of initial encounter complexes. It is therefore not clear whether the protein-protein binding interfaces observed in the ternary complexes are still preferred without PROTACs. Based on the definition, the positive measured values of binding cooperativity indicate that protein-protein interactions are energetically favorable to drive the formation of corresponding ternary complexes. As a result, the protein-protein binding interfaces in these complexes should be more likely adopted than other potential interfaces. In contrast, the negative measured values of binding cooperativity indicate that protein-protein interactions are energetically unfavorable during the process of forming corresponding ternary complexes, suggesting that without PROTACs, other alternatives binding interfaces could be adopted more easily than the interfaces existing in current complexes.

In order to test these assumptions, we artificially built decoy structures of protein-protein interactions for the ternary complex that has positive binding cooperativity. The decoy structures were constructed using molecular docking by HDOCK. A more detailed description of HDOCK can be found in the **Methods**. Specifically, the complex formed between the target protein BRD4^BD2^ and the ligase VHL (PDB id 5T35) was used as a test system. The monomeric structure of BRD4^BD2^ (PDB id 5UEU) and VHL (PDB id 4W9H) were used as the inputs of HDOCK. The top 10 models generated from HDOCK for the complex were then selected. Additionally, considering that the binding interfaces in these models were not as refined as in the experimentally derived structure, we did not compare binding interfaces in decoy structures to the native interface in 5T35. Instead, we compared them to an artificially modeled interface by docking the monomeric BRD4^BD2^ and VHL to the native complex, so that the comparison could be done on a more equitable basis. **Figure 3a** shows the structures of all decoy and native-like complex. The target proteins in these complexes were superimposed together, as colored in red. The ligase with the native-like binding interface is colored in green, while the ligases in all the decoy complexes are colored in blue. A large diversity of binding interfaces is observed in the figure.

**Figure 3:**
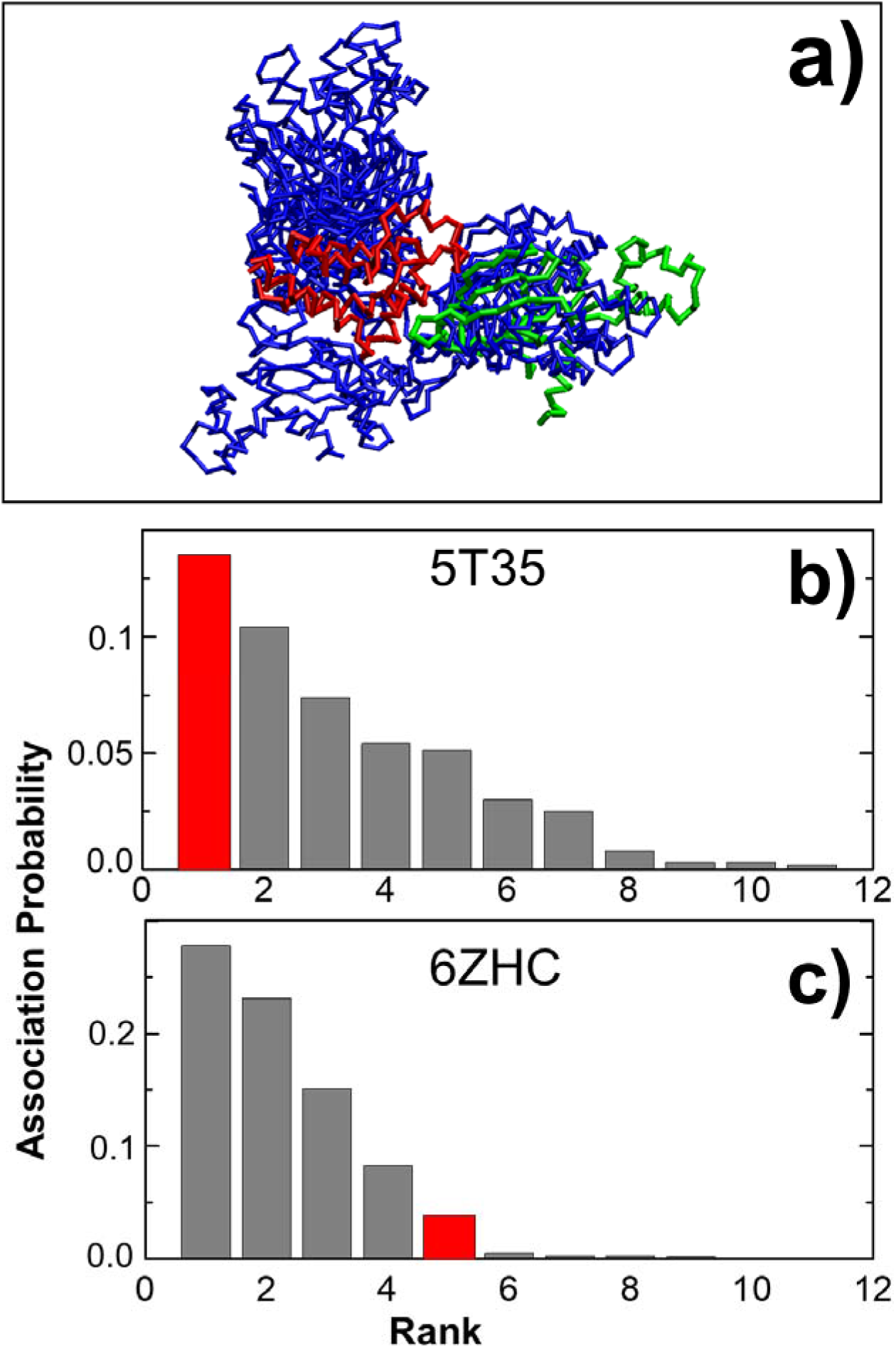
We compared the decoy structures of the protein complex (PDB id 5T35) generated by molecular docking with its native conformation **(a)**. The target proteins are colored in red and were superimposed together, while the ligases in the native and decoy complexes are colored in green and blue, respectively. KMC were used to calculate the association probability for all complexes that formed through either decoy or native-like interface. They are plotted as a histogram in **(b)** in decreasing order. The figure shows that the native complex (red bar) has the highest association probability than all the decoy complexes (grey bars). On the other hand, another complex with measured negative binding cooperativity (PDB id 6ZHC) was selected as a control study. The comparison of calculated association probabilities between native and decoy structures is plotted in **(c)** for this complex. The figure shows that many decoy complexes have higher association probabilities than the native complex.

For each complex that formed through either decoy or native-like interface, the KMC was apply to generate 10^3^ simulation trajectories from different initial configurations, in which BRD4^BD2^ and VHL were separated into monomers. Among these trajectories, we counted the number of trajectories in which complexes were formed through the corresponding interface. We further calculated the association probability of forming the 10 decoy complexes, as well as the probability of forming the interaction as observed in the ternary complex. The final calculated association probabilities of interaction between BRD4^BD2^ and VHL through native-like and all 10 decoy interfaces are plotted as a histogram in **Figure 3b**, ranked in decreasing order. In the figure, the probability of protein-protein association through the native-like interface is shown as the red bar, while the probabilities of association through the decoy interfaces are shown as the grey bars. Our calculated probability is 0.135 for the association through the native-like interface. Although it is lower than 0.338, where the experimental structure of the binding interface (PDB id 5T35) was used, the probability is still higher than all the others that were formed through decoy interfaces. This result indicates that the protein-protein interfaces in the ternary complexes that have positive cooperativity is highly preferred over other potential binding modes. In these cases, the protein-protein interactions play a positive role in driving the formation of ternary complexes.

In a control study, we artificially built decoy structures of protein-protein interactions formed between the target protein BCL2L and the ligase VHL (PDB id 6ZHC) by HDOCK. The ternary complex in this case has a negative binding cooperativity. The top 10 models of the protein complex were selected. Following the same procedure, we applied the KMC to generate 10^3^ simulation trajectories for these 10 decoy complexes. Each trajectory was started from a different initial configuration. We then counted the number of trajectories in which complexes were formed through the corresponding interface at the end of all trajectories. The association probability of forming these 10 decoy complexes were thus calculated. The probabilities of interaction between BCL2L and VHL through decoy interfaces are compared with the probability of interaction through the native interface. These probabilities are plotted as a histogram in **Figure 3c**, ranked in decreasing order. The figure shows that, different from the ternary complex with positive cooperativity, the probability of protein-protein association through the native interface (red bar) is not ranked higher than some of those through decoy interfaces (grey bars). More specifically, among the 10 decoy interfaces, four of them have a higher association probability than the native interface. This result indicates some alternative protein-protein binding modes might be energetically more favorable than the native conformation of the ternary complex with negative cooperativity.

In summary, we compared the protein-protein association through different interfaces between ternary complex with positive and negative cooperativities. The native-like binding interface in the complex with positive cooperativity shows a higher association probability than all the other alternative interfaces. Protein-protein interactions play a dominant role in driving the formation of these ternary complexes. In contrast, the native-like binding interface in the complex with negative cooperativity could be less favored than some other alternative interfaces. We speculate that the design of PROTACs that target these interfaces might lead to new ternary complexes with a higher value of binding cooperativity.

### Target proteins with high structural similarity could bind to their ligase quite differently: a case study of CDK4/6

Because cyclin-dependent kinases 4 and 6 (CDK4/6) play an essential role in regulating the cell-cycle transition from G1 to S-phase, they are promising targets for cancer therapy [28]. Although the structures of CDK4 and CDK6 are highly similar, they have diverse functions in cells [29]. The development of therapeutics that can selectively inhibit specific kinases therefore is highly desirable. Unfortunately, this selectivity cannot be easily achieved due to the fact that current CDK4/6 inhibitors all target their conserved ATP-binding pockets. This offers the opportunity to other alternative strategies. For instance, bringing different kinases to ligase for degradation might reach this goal. Previous experiments have shown the feasibility of targeting specific CDK through the design of various PROTAC-based molecules. We assume that this selectivity is driven by the difference in the interactions between the two proteins and ligase.

In order to test this assumption, we applied KMC to simulate the association between ligase and CDK4/CDK6, and explored the differences in these two systems. PROTAC-based degraders have been developed by linking a E3 ligase (CRBN) binding moiety, such as thalidomide, directly to small-molecule inhibitors, such as palbociclib and ribociclib [30]. The structural evidences show that the aminopyrimidine moiety of these inhibitors form hydrogen bonds with the backbone of the kinases, while their piperazine rings directly interact with a solvent-exposed ridge of the proteins [31]. Unfortunately, because the structure of ternary complexes consisting of CDK4/CDK6 and CRBN is unavailable, their association was modeled computationally as follows. In order to minimize the potential biases, the structures of both kinases were built by AlphaFold2 [32] from their sequences. The sequence of CDK4 was adopted from the UniProt ID P11802. The sequence of CDK6 was adopted from the UniProt ID Q00534. On the other hand, the structure of CRBN was adopted from the PDB ID 6BN7. We then placed the monomeric kinase and ligase separately in a simulation box with random orientation. Starting from this initial configuration, two proteins were moved against each other by KMC algorithm.

For both CDK4 and CDK6 systems, the KMC generated 10^3^ simulation trajectories from different initial configurations. Within each trajectory, we calculated the distance between the thalidomide binding pocket in CRBN and the small-molecule inhibitor binding pocket in the kinases. The changes of this calculated distance along the simulation time are shown in **Figure 4a** for some representative trajectories. As the ligase and the kinase were initially separated in the simulation box, the distance between two binding pockets was large at the beginning of the trajectories. The figure shows that, in some cases, the ligase and the kinase cannot form a complex by the end of the simulation trajectory represented by the black curve, in which the distance between their binding pockets remained large with high fluctuations. In some other cases, however, the distance between the thalidomide binding pocket in CRBN and the small-molecule inhibitor binding pocket in CDK4 or CDK6 reduced to a small value before the end of the trajectory, as shown by the red curve in the figure. This indicates that the interactions between ligase and kinase could guide their diffusions, bring their corresponding binding pockets close to each other, and lead to the formation of a ternary complex. The initial and final configurations of the trajectory are plotted in **Figure 4b** and **Figure 4c**, respectively.

**Figure 4:**
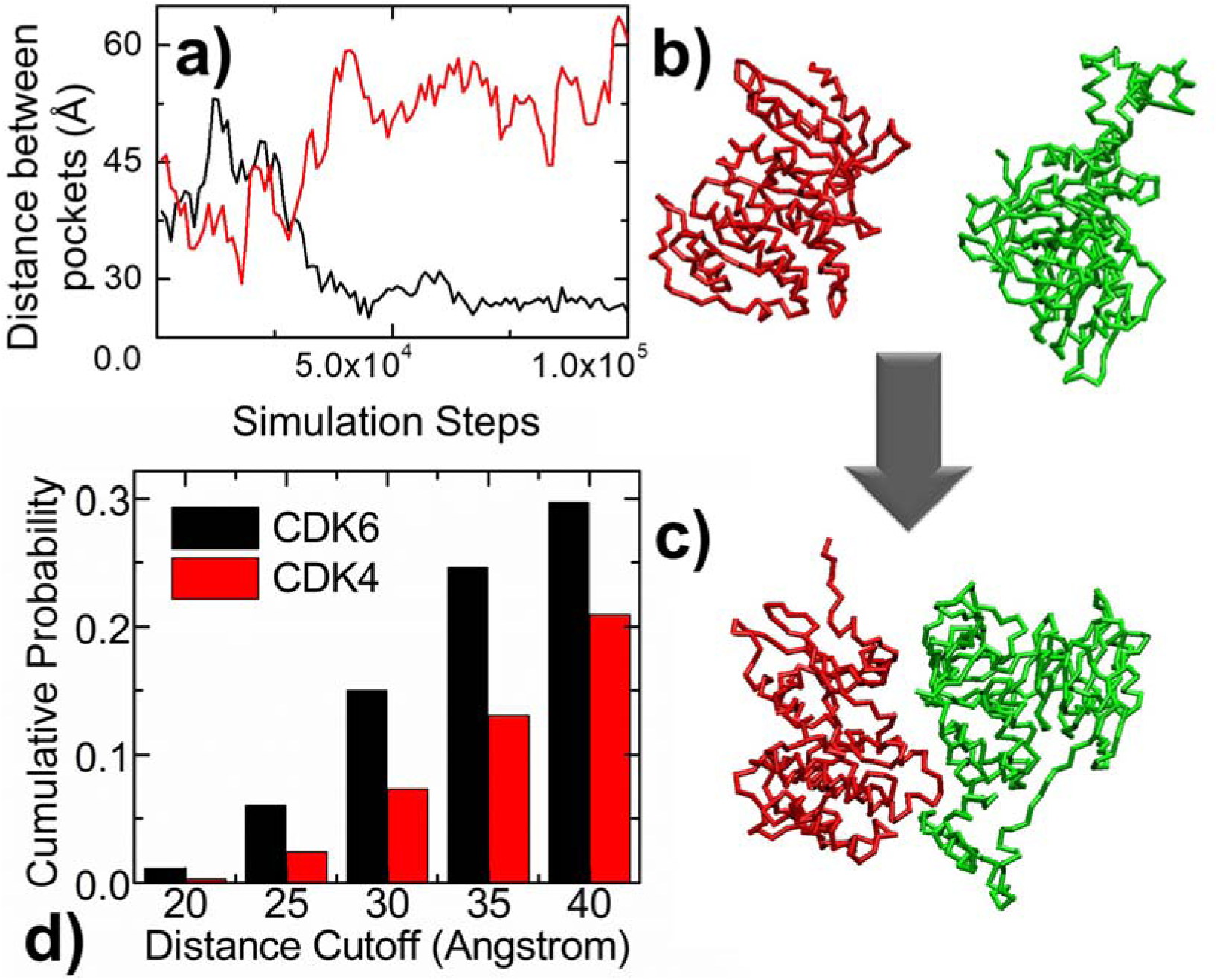
We applied KMC simulation to generate 10^3^ simulation trajectories from different initial configurations for both CDK4 and CDK6 systems. We calculated the distance between the thalidomide binding pocket in CRBN and the small-molecule inhibitor binding pocket in CDK4/CDK6 along the simulation, as shown in **(a)** for two representative trajectories. The black curve in the figure indicates the ligase and the kinase cannot form a complex by the end of the simulation, while the red curve indicates the interactions between ligase and kinase bring their corresponding binding pockets close to each other and lead to the formation of a ternary complex. The initial and final configurations of the trajectory are plotted in **(b)** and **(c)**, respectively. Finally, the statistical comparison between CDK4 and CDK6 among all 10^3^ simulation trajectories is shown by the histogram in **(d)**.

The statistical comparison between CDK4 and CDK6 among all 10^3^ simulation trajectories is shown by the histogram in **Figure 4d**. The x axis of the histogram denotes the cutoff values of the distance between the binding pocket in the kinase and the binding pocket in the ligase. The y axis gives the probability of finding the trajectories in which the pocket distance between the ligase and the kinase was below the corresponding cutoff. Although CDK4 and CDK6 share high structural similarity, the figure shows that they interact differently with CRBN. In detail, while forming complexes, the binding pocket of CDK6 was much more frequently presented near the binding pocket of CRBN than CDK4. For instance, among the 1000 trajectories of the CDK6 system, 60 of them have pocket distances below the cutoff of 25Å. On the other hand, only 24 trajectories of the CDK4 system have pocket distances below the same cutoff.

A previous study showed that adding an extended PEG-3 linker to an original ribocicilib-based dual CDK4/6 degrader can make it become a selective CDK6 degrader. Because both ligase and kinase binding moieties in the degrader remained unchanged, it suggests that the length of the linker leads to the selectivity. More specifically, a longer and more flexible linker made the degrader only target CDK6 but not CDK4. Our simulation results here indicate that the binding pockets in CRBN and CDK6 are more likely to be close to each other during their association than the binding pockets in CRBN and CDK4. Consequently, degraders have higher probabilities of being engaged in their binding pockets, leading to the formation of ternary complexes. This could explain the experimental observation that PROTACs with extended linkers can still be effective against CDK6, but not against CDK4.

In summary, KMC was applied to two related but functionally diverse proteins. We found that their interactions with the ligase are not as similar as their structures. Our study provided insights into the design of PROTAC that can selectively target specific proteins.

## Conclusions

Not all functional proteins can be targeted pharmacologically by traditional small-molecule inhibitors. The idea of targeted protein degradation tremendously expands the druggable space. PROTACs, as a well-developed example of targeted protein degradation, link a target protein to the E3 ligase, leading to the formation of a ternary complex. The properties of protein-protein interactions are crucial to the productivity of PROTAC-induced ternary complexes. It was believed that an energetically favored binding interface between proteins results in positive cooperativity during ternary complex formation, further ensures selectivity, and elicits potent degradation. The impacts of protein-protein interactions on PROTAC-based targeted degradation, however, have not been systematically tested by computational simulations. In this paper, kinetic Monte-Carlo simulation has been applied to a most updated benchmark set of non-redundant PROTAC-based ternary complexes. We found that proteins in the complexes of the benchmark are more likely to interact with each other if their measured binding cooperativities are positive. Moreover, we compared the protein-protein association through different interfaces generated from molecular docking. The native-like binding interface shows a higher association probability than all the other alternative interfaces only in the complex with positive cooperativity. These results support the idea that the formation of ternary complexes is closely regulated by the binary interactions between proteins. In the last test, by comparing how CDK4 and CDK6 interact with the ligase, we found that CDK6’s binding pocket appeared closely to the binding pocket of CRBN, much more frequently than CDK4, in spite of their high structural similarity. Altogether, our study paves the way for understanding the role of protein-protein interactions in PROTACE-induced ternary complex formation. It can potentially help in searching for degraders that selectively target specific proteins.

## Methods

### Constructing the benchmark dataset of PROTAC-based ternary complexes

In order to build the dataset, we first searched the PDB using the keyword “PROTAC”. This gave us a total number of 113 entries. A large portion of these entries contain the structures of only target proteins or the structures of only ligases. We only considered the structures of ternary complexes, which should contain both target proteins and ligases. After the removal of all binary complexes, 46 entries were left. However, there are redundancies among these remaining entries. Certain ternary complexes with the same target protein and ligase are over-represented in the PDB database. For example, both PDB IDs 5T35 and 6SIS represent the structure of the ternary complex consisting of the target protein BRD4^BD2^ and the ligase VHL. We manually check this redundancy. For entries representing the same ternary complex, only one was kept in our dataset. As a result, 9 ternary complexes were finally selected.

For all these 9 selected ternary complexes, we downloaded their atomic coordinates from the PDB as the inputs of our KMC simulations. We further obtained the value of their experimentally measured binding cooperativity from the literature. Detailed information of our collected dataset is summarized in **Table 1**. The first column of the table indicates the PDB ID of each ternary complex in the dataset. The second and third columns denote the names of the corresponding target proteins and ligases, respectively. The fifth column shows the experimentally measured binding cooperativity of each ternary complex. The positive cooperativity was marked as “+”, while the negative cooperativity was marked as “-”. Their corresponding references are cited in the last column of the table. Finally, the association probabilities calculated from the KMC simulations are listed in the fourth column for all ternary complexes.

### Generating decoys of protein complexes by protein-protein docking

Giving the structures of a ligase and a target protein as inputs, the HDOCK server was applied to perform global docking between the two proteins [33]. The template-free docking mode was adopted in order to generate diversified binding interfaces. The server first uses a fast Fourier transform (FFT)–based global search method to sample putative binding interfaces between the two query proteins. The sampled binding interfaces are then evaluated by a knowledge-based scoring function learned from the available structures of protein–protein interactions. The performance of this hierarchical docking algorithm has been demonstrated by the community-wide experiment, Critical Assessment of PRediction of Interactions (CAPRI) [34]. Finally, the top 10 models of protein complexes were selected and their atomic coordinates were downloaded from the server.

### Simulating the protein-protein association by coarse-grained kinetic Monte-Carlo algorithm

We applied our previously developed kinetic Monte-Carlo simulation method to study the association between target proteins and their corresponding ligases. In the simulation, protein structures are described by a coarse-grained model. Each residue of a protein is simplified by two points, one is its Cα atom and the other represents the side-chain. A target protein and its ligase were separated and randomly placed in a three-dimensional simulation box as the initial configuration. Following the initial configuration, random translational and rotational movements were carried out for both proteins within each simulation step. The amplitudes of these movements are determined by the diffusion constant of each protein. A physics-based scoring function was then used to guide diffusions. The scoring function contains two terms to describe the electrostatic and hydrophobic interactions, as well as an additional penalty to avoid clashes between proteins. Based on the calculated scoring function, Metropolis criterion was applied to determine whether the diffusional movements were accepted or not. This process was iterated until an encounter complex was formed between the target protein and ligase through their corresponding binding interface. Otherwise, if two proteins could not form interactions, the simulation ended after it reached the maximal time duration. In order to effectively estimate the association probability for a given ternary complex, a large number of simulation trajectories were performed under different initial configurations. Two proteins could successfully form complexes in some trajectories, but diffuse far away from each other in others. The association probability was derived by counting the number of trajectories in which complexes were successfully formed over the total number of trajectories.

## Data and code availability

All data from this study, including the benchmark of ternary complexes, the relevant source codes of the kinetic Monte-Carlo simulation and the results generated from the simulations can be found in the GitHub repository https://github.com/wulab-github/PROTAC_KMC.

## Acknowledgement

This work was supported by the National Institutes of Health under Grant Numbers R01GM120238 and R01GM122804. The work is also partially supported by a start-up grant from Albert Einstein College of Medicine. Computational support was provided by Albert Einstein College of Medicine High Performance Computing Center.

## Author Contributions

Z.S., S.Y. and Y.W. designed research; Z. S. and Y.W. performed research; Z.S. S.Y. and Y.W. analyzed data; S.Y. and Y.W. wrote the paper.

## Additional Information

### Competing financial interests

The authors declare no competing financial interests.

## Notes

### Competing Interest Statement

The authors have declared no competing interest.

